# Measuring PETase enzyme kinetics by single-molecule microscopy

**DOI:** 10.1101/2024.04.24.590935

**Authors:** Yuwei Zhang, William O. Hancock

## Abstract

Polyethylene terephthalate (PET) is one of the most widely produced man-made polymers and is a significant contributor to microplastics pollution. The environmental and human health impacts of microplastics pollution have motivated a concerted effort to develop microbe- and enzyme-based strategies to degrade PET and similar plastics. A PETase derived from the bacteria *Ideonella sakaiensis* was previously shown to enzymatically degrade PET, triggering multidisciplinary efforts to improve the robustness and activity of this and other PETases. However, because these enzymes only erode the surface of the insoluble PET substrate, it is difficult to measure standard kinetic parameters, such as k_on_, k_off_ and k_cat_, complicating interpretation of the activity of mutants using traditional enzyme kinetics frameworks. To address this challenge, we developed a single-molecule microscopy assay that quantifies the landing rate and binding duration of quantum dot-labeled PETase enzymes interacting with a surface-immobilized PET film. Wild-type PETase binding durations were well fit by a biexponential with a fast population having a 2.7 s time constant, interpreted as active binding events, and a slow population interpreted as non-specific binding interactions that last tens of seconds. A previously described hyperactive mutant, S238F/W159H had both a faster on-rate and a slower off-rate than wild-type PETase, potentially explaining its enhanced activity. Because this single-molecule approach provides a more detailed mechanistic picture of PETase enzymatic activity than standard bulk assays, it should aid future efforts to engineer more robust and active PETases to combat global microplastics pollution.

**Statement of significance:** Plastic pollution is a global environmental and human health problem. PETases are recently discovered enzymes that degrade the ubiquitous plastic polyethylene terephthalate (PET). A push is underway to understand and optimize these enzymes to enable large-scale microplastics remediation. Here, we use single-molecule fluorescence microscopy to visualize the interactions of PETase enzyme molecules with a thin film of PET. We identify specific binding interactions of a few seconds that differ between wild-type and PETase mutants that have been previously shown to have altered activities. These single-molecule investigations provide a new window into the mechanism and activity of PETase enzymes, and provide a platform for characterizing and optimizing novel PETases with improved function and stability.

## Introduction

Polyethylene terephthalate (PET) is one of the most widely produced man-made polymers, and is derived from petroleum by the esterification of terephthalic acid and ethylene glycol. Accumulation of PET and other microplastics in the environment is a pressing global problem that has motivated efforts to develop microbe- and enzyme-based strategies to degrade PET (1, 2). In 2016, Yoshida *et al*. reported a newly discovered bacterium, *Ideonella sakaiensis* 201-F6, that was able to use PET as its major carbon and energy source for growth. The PETase enzyme derived from this microbe, *Is*PETase, was shown to convert PET to mono(2-hydroxyethyl) terephthalic acid (MHET), with trace amounts of terephthalic acid (TPA) and bis(2-hydroxyethyl)-TPA (BHET) as secondary products (3). This discovery set off a global interdisciplinary effort to identify, characterize and optimize PET degrading enzymes, with the hope that these biodegradation approaches can be scaled up to reduce microplastics pollution in the environment (4, 5).

Current methods to evaluate the specific activity of PETases have included analysis of PET surface erosion by scanning electron microscope (SEM), quantifying decreases in PET crystallinity using differential scanning calorimetry, and measuring product released by HPLC (6–8). These bulk assays provide quantitative information on enzyme activity, but relating these measurements to the specific enzyme kinetic parameters of the PETase enzymes is complicated by the insolubility of PET and the fact that enzymes only act on exposed surfaces. Alternatively, there are turbidimetric or colorimetric methods for PETase enzyme activity, such as the soluble substrates p-nitrophenyl acetate (pNPA) and p-nitrophenyl butyrate (pNPB), but these soluble substrates are poor analogs for crystalline PET (7, 9).

Like PETases, cellulases and cutinases are enzymes that react only with the surface of their insoluble subtrates (10). Because the effective surface: volume relationship is generally not known, it is difficult to directly apply the Michaelis-Menten formalism to these reactions. Specifically, the resulting K_M_ values are not in typical molar concentrations, making it difficult to extract the bimolecular on- and off-rates, k_on_ and k_off_, as well as the enzyme turnover rate, k_cat_. One approach that has been used extensively on enzymes like cellulases or cutinases is to vary the enzyme concentration rather than the substrate concentration in the solution, such that enzymes partition to bound and soluble fractions (11–13). However, even here the available area, *A*_0_, is a key parameter that must be estimated rather than precisely measured. In contrast, single-molecule approaches directly measure binding rates and durations, and thus they offer more direct access to true biochemical parameters of enzymes. Further, because the activity of the enzyme (rather than the substrate) is monitored, they sidestep the requirement of relating exposed surface area to substrate mass or volume, which bulk assays typically require. To this end, by imaging the cellulase Cel7A binding to and moving along its insoluble substrate, cellulose, we were able to extract enzyme kinetic parameters from the single-molecule tracking dynamics (14).

Previous work that used bulk enzymatic assays and molecular dynamics simulations to investigate key residues in the substrate binding cleft of *Ideonella sakaiensis* PETase identified two mutants with altered activities (6). A hyperactive mutant, S238F/W159H, was created that narrowed the substrate binding cleft towards that of native cutinase, and an impaired mutant, W185A was created by mutating a highly mobile tryptophan thought to play a role in substrate binding. The altered enzyme activities were measured by imaging the surface erosion of PET using electron microscopy and by monitoring changes in the crystallinity of the PET substrate. These experimental approaches clearly showed changes in enzyme activity, particularly the enhanced activity of the hyperactive mutant, but because the methodology did not quantify k_on_, k_off_ and k_cat_, it is difficult to define in enzymatic terms which transition rates are altered by the mutations.

In the present work, we measured the single-molecule binding dynamics of *Ideonella sakaiensis* PETase wild-type and both the hyperactive and impaired mutants characterized previously. We used total internal reflection fluorescence microscope (TIRFM) of quantum dot (Qdot) labeled PETase reversibly binding to glass coverslips functionalized with a thin film of PET. By measuring the residence time of the enzymes on the substrate, we calculate an apparent off-rate that provides information about both the turnover rate of the enzyme and its binding affinity to the PET substrate. This work provides new insights into PETase function and introduces a new approach for investigating the enzymology of other enzymes that react with insoluble substrates.

## Materials and methods

### Expression and purification of PETase enzymes

DNA plasmids for wild-type PETase (PETase, *I. sakaiensis*, accession number: A0A0K8P6T7, Addgene number: #112202), hyperactive mutant (Addgene number: #112203) and the impaired mutant (Addgene number: #112204) were purchased from Addgene (6). Plasmids were transformed into *E. coli* C41(DE3) (Lucigen) and inoculated on a Nutrient Agar plate containing 100 μg/mL ampicillin (VWR; Cas: 69-52-3). After culturing overnight at 37°C, single colonies for each variant were inoculated into a started culture of 5 mL Terrific Broth media (Thermo Scientific) containing 100 μg/mL ampicillin and grown at 37°C overnight in a 180 rpm shaking incubator. Each starter culture was then transferred into 2 L flasks with 400 mL media containing 100 μg/mL ampicillin and grown at 37°C at 200 rpm until the optical density at 600 nm reached 0.6-0.8. Protein expression was induced by adding 1 mM isopropyl β-D-1-thiogalactopyranoside (IPTG) (EMD Millipore Corp.; Cas: 367-93-1). Cells were then incubated for 18 to 20 hours at 20°C at 200 rpm, harvested by centrifugation at 5000 rpm for 20 min, and stored at -80°C for later purification. Frozen cell pellets were resuspended in a lysis buffer (300 mM NaCl, 10 mM imidazole, 20 mM Tris HCl, pH 7.4,) and lysed by sonication. Lysate was clarified by centrifugation at 45,000 rpm for 35 minutes. The supernatant was then applied to a 2 mL Ni Sepharose 6 Fast Flow (Cytiva) column and eluted by elution buffer containing 300 mM NaCl, 300 mM imidazole, 20 mM Tris HCl, pH 7.5.

### Microscopy and image analysis

PET film (Goodfellow, UK) was cleaned in purified water by sonication, wiped with 100% ethanol, cut into pieces, and dissolved in trifluoroacetic acid (TFA) (Sigma Aldrich; Cas: 76-05-1) to a concentration of 1 wt%. 5-7 μL of PET solution was dripped onto a plasma cleaned coverslip, covered with a second cleaned coverslip, and the two coverslips were then separated and dried in air at room temperature for 30 min. Next, the coverslips were transferred to a 120°C oven for 20 min to achieve a smooth PET coating, followed by cooling at room temperature. Flow cells were assembled with the PET-coated coverslips using double-sided tape. The PET surface was blocked to minimize nonspecific protein adsorption by flowing 0.33 mg/mL casein into the flow cell for 3 min, followed by an enzyme solution consisting of 8 nM wild-type PETase or its mutant labeled with 0.8 nM Qdot_655_ (Thermo Scientific), 2 mM 2-Mercaptoethanol (Sigma-Aldrich; Cas: 60-24-2), and 4 mM 4-Nitrophenyl acetate (Sigma-Aldrich; Cas: 830-03-5) in a buffer containing 100 mM NaCl and 50 mM sodium phosphate, pH 7.4. The Qdot_655_-labeled PETase was imaged by total internal reflection fluorescence microscopy (TIRFM) using a 488 nm laser (70 mW or 150 mW) on a custom-built microscope (15). Movies began immediately before the enzyme solution was added to the flow cells. All experiments were performed at 22°C. The 10:1 enzyme:Qdot ratio was chosen because binding events were quite rare and lower ratios resulted in insufficient particle counts to accurately calculate landing rate and duration parameters.

PETase landing events and binding durations on the surface-immobilized PET film were measured by Kymograph analysis in ImageJ, as follows. Image stacks consist of a cube of x-y images at sequential times, t. First, the “Reslice” command was used to generate multiple x-t (horizontal) slices at varying y (vertical). The “Output spacing” was set to 5 pixels, which corresponds to the ∼300 nm full width at half maximum of the point spread function of our microscope (15). Next, the “Grouped z projection” command was used to compress the multiple slices along the y axis into a single x-t image (see Fig. 1). From this kymograph, the binding durations are measured by the length of the vertical lines, where one pixel equals one frame, and the landing rate is measured by counting the total number of events throughout the entire movie. This approach of generating an x-t Kymograph is similar in principle to the approach of Pettersson *et al* (16). Binding duration distributions were fit to biexponentials in Python 3.9.19 using the ‘scipy.optimize’ library.

**Figure 1.**
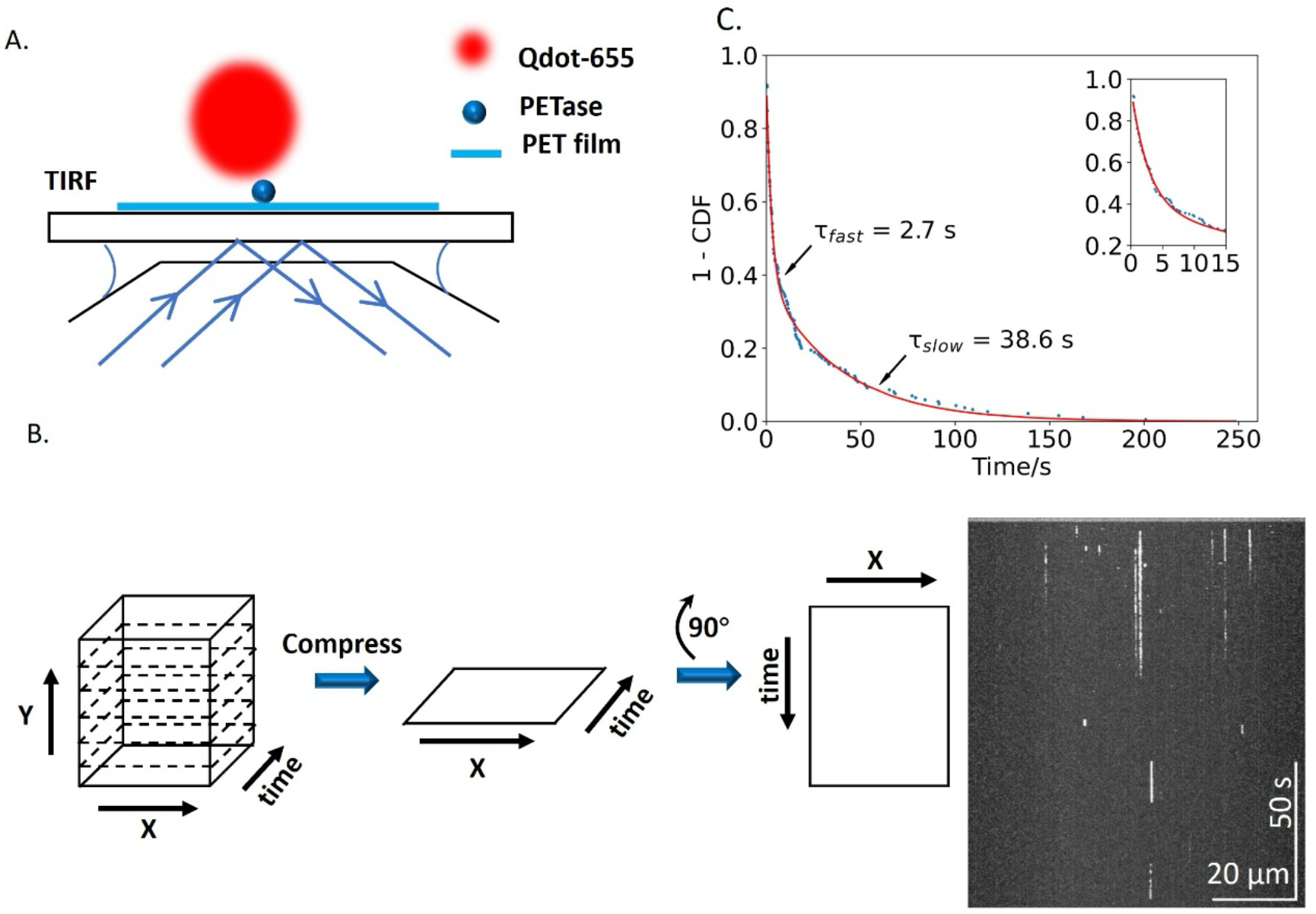
PETase binds reversibly to PET film. (A) Experimental design for single-molecule TIRF analysis of Qdot-labeled PETase interacting with surface-immobilized PET film. Image is not to scale. (B) Kymograph analysis. A movie, consisting of a stack of x-y images at various t is captured and resliced into x-t frames for each y, and then compressed into a single x-t image. The image is then rotated by 90° to generate a single x-t kymograph (shown at right), where vertical streaks are single-molecule binding events of varying durations. (C) Binding durations of wild-type PETase, plotted as 1 – cumulative distribution function (1-cdf). Fitting to a biexponential function reveals a fast phase, representing reversible binding interactions, as well as a slow phase. Inset: first 15 s to show details of fast phase.

## Results

### Qdot_655_-labeled PETase binds reversibly to PET substrate

To observe the behavior of PETase enzymes on an immobilized PET surface, PET was dissolved in trifluoroacetic acid and dripped onto the surface of a coverslip, which was then covered by another plasma cleaned coverslip to make thin PET film on the surface of both coverslips. The PET-functionalized coverslip was incorporated into a flow cell, and 0.33 mg/ml casein was flowed through to reduce nonspecific protein adsorption. *Ideonella sakaiensis* PETase (Addgene Number: #112202) (6) was bacterially expressed and biotinylated (see Methods), and combined with streptavidin-coated Qdot particles at a 10:1 enzyme:Qdot ratio. The PETase-Qdot complexes were then introduced into the flow cell and allowed to interact with the PET surface. Landing events were imaged by total internal reflection fluorescence microscopy (TIRFM) on a custom-built multimodal microscope (15), and movies were obtained of the enzymes reversibly binding to the surface-immobilized PET.

To quantify the reversible binding interactions, we developed a kymograph-based approach (Fig. 1B; described in Methods) to measure the frequency and duration of PETase binding events. The 5 frames/s video consists of a stack of x-y images through time. The binding events in our assay are focal-limited spots of diameter ∼300 nm, corresponding to the point-spread function of the microscope (15). Because the events are relatively rare, we can collapse a given image in y-without superimposing different events. When this image is rotated by 90°, the resulting image corresponds to the positions in x along the horizontal axis and time along the vertical axis (Fig. 1B). From this kymograph, the binding durations are measured by the length of the vertical lines, where one pixel equals one frame, and the landing rate is measured by counting the total number of events throughout the entire movie.

We found that the distribution of binding durations of wild-type PETase was well fit by a double exponential function consisting of a fast phase with a time constant of 2.7 ± 0.12 s (fit ± SE of the fit for all time constants) and a slow phase with a time constant of 38.6 s (Fig. 1C).

### Slow binding phase is consistent with non-specific binding

To better understand the origin of the fast and slow binding populations, we carried out a control experiment using Qdots with no PETase bound. Using 70 mW laser power, Qdots were observed to reversibly bind to the surface, and the binding durations followed a biexponential with a fast phase of 1.5 ± 0.08 s (faster than the PETase fast phase of 2.7 s) and a slow phase of 51.1 ± 2.0 s (Fig. 2A). We hypothesized that the slow time constant may be due to nonspecific and/or irreversible binding of Qdots to the surface, with the apparent unbinding rate being due to photobleaching or blinking of the Qdots. If this were the case, then increasing laser power to accelerate bleaching should shorten the time constant of the slow phase. To test this, we repeated the control Qdot experiment using a 150 mW laser illumination and found that the slow time constant shortened from 51.1 s to 23.3 s, as expected, while the fast time constant remained at 1.5 s (Fig. 2B). This result is consistent with the slow phase being due to irreversible binding events whose durations are truncated by photobleaching or blinking, rather than dissociation.

**Figure 2.**
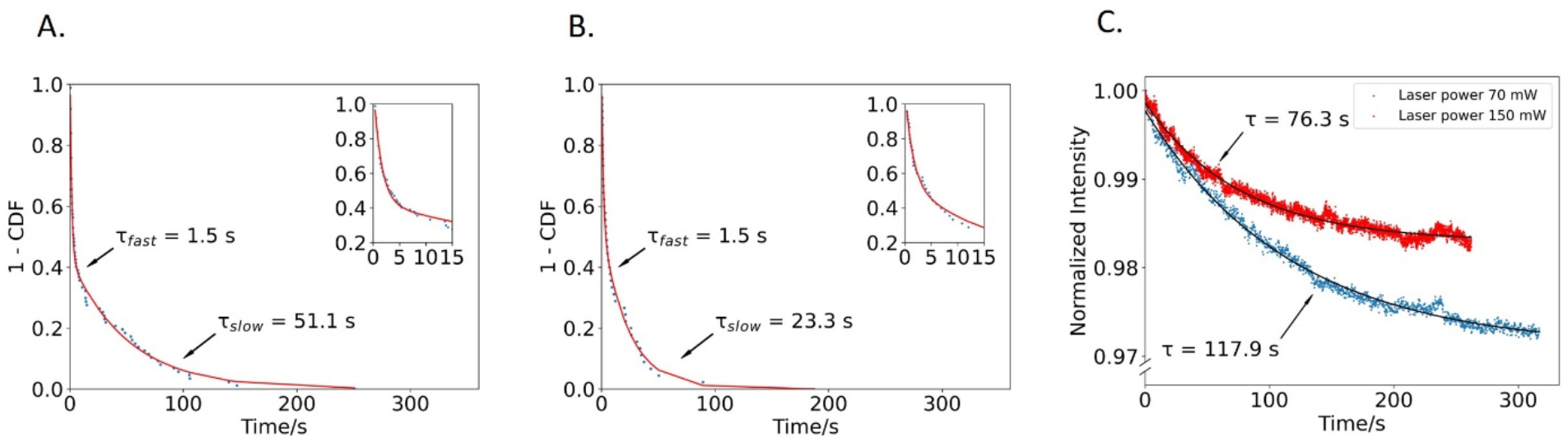
Time constant of the slow phase varies with laser power. (A) Binding durations of PETase-free Qdots at 70 mW illumination with biexponential fit. Inset: early times to show fast phase. (B) Binding durations of PETase-free Qdots at 150 mW illumination with biexponential fit. (C) Photobleaching rates of immobilized Qdots at varying illumination intensities. Particles were nonspecifically bound to the surface and the average image intensity over time plotted at 70 mW and 150 mW illumination intensities. Fit to exponential fall gives time constants for Qdot photobleaching of 117.9 s and 76.3 s at 70 mW and 150 mW illumination, respectively.

To further validate our photobleaching hypothesis, we immobilized Qdots on the surface and quantified their population-level photobleaching rate. To do this, we flowed 50 pM Qdots (lacking PETase) into the flow cell without first blocking the PET surface to prevent nonspecific adsorption, washed away any free Qdots that did not bind, and imaged these nonspecifically-bound Qdots under continuous laser illumination. The average fluorescence intensity of the image fell exponentially over time due to photobleaching of the immobilized Qdots (Fig. 2C). At 70 mW laser power, the time constant was 118 s, and at 150 mW laser power, the time constant fell to 76.3 s, consistent with faster Qdot photobleaching at higher laser powers. This population-level result reinforced our single-molecule result in Fig. 2AB, supporting our conclusion that the slow time constant in the PETase experiments (Fig. 1) is due to slow photobleaching of Qdots that are nonspecifically bound to the surface.

### On- and off-rates of hyperactive and impaired PETases differ from wild-type

We next used our single-molecule PETase assay to investigate the reversible binding dynamics of two previously described PETase mutants. Austin *et al*. generated a hyperactive double mutant, S238F/W159H, which was designed to narrow the active site of the enzyme and was found to degrade crystalline PET more effectively than wild-type (6). They also generated an inactive mutant W185A, which was found to have impaired activity compared to wild-type PETase (6).

Repeating our wild-type approach, we bacterially expressed, purified, and biotinylated the two mutants, adsorbed them to Qdots, and quantified their binding durations. Comparing the fast phases, which correspond to reversible binding, the hyperactive mutant had a time constant of 4.0 ± 0.07 s (fit ± SE of the fit), longer than the wild-type (2.7 ± 0.12 s), whereas the impaired variant had a time constant of 1.7 ± 0.18 s, shorter than wild-type (Fig. 3A-C). These binding durations are consistent with the hyperactive mutant having a higher PET binding affinity, and the impaired mutant having a lower binding affinity than wild-type.

**Figure 3.**
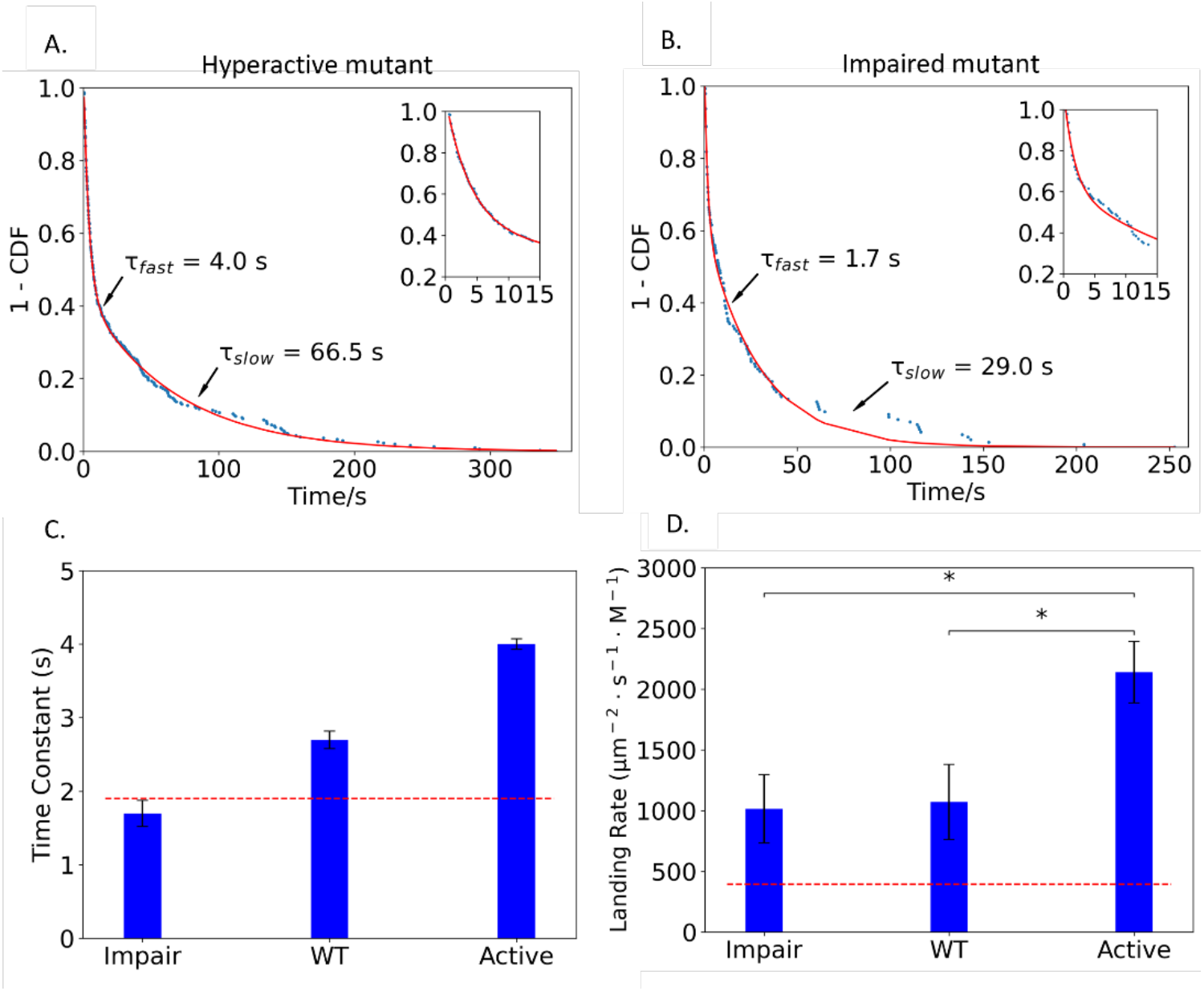
Hyperactive and impaired mutants have different fast-phase time constants and overall landing rates. (A) Distribution of binding durations of hyperactive PETase mutant, fit by a biexponential. Inset: first 15 s to show details of fast phase. (B) Distribution of binding durations of impaired PETase mutant, fit by biexponential. Inset: first 15 s to show details of fast phase. (C) Time constants of fast phase for wild-type (from Fig. 1C), and impaired and hyperactive mutant PETase (From Fig. 3A and B, respectively). Dashed line is time constant fast phase from Qdot control (see Fig. S1 for Qdot binding duration distribution). Error bars are standard error of the fit. (D) Single-molecule PETase landing rates, including all events independent of duration. Dashed line indicates PETase-free Qdot control landing rate. Error bars are SEM with N = 7 to 11 screens. Asterisks denote data are significantly different from one another with p <0.05 by a two-sample t-test. Additionally, all landing rates were significantly different from the Qdot control (red dashed line), with p<0.05 by two sample t-test.

To quantify the relative on-rates for PET binding, we quantified the binding frequency of the three PETase enzymes along with the Qdot control (Fig. 3D). Binding rates were normalized to the enzyme concentration and to the area of the microscope image. The binding rate of the impaired mutant, 1014 ± 282 μm^-2^ · s^-1^ · M^-1^ (mean ± SEM for N=7 movies) was similar to the wild-type rate of 1070 ± 310 μm^-2^ · s^-1^ · M^-1^ (N = 9). Notably, the hyperactive mutant landed more frequently, at 2141 ± 254 μm^-2^ · s^-1^ · M^-1^ (N = 7) than wild-type. Taken together, the results suggest that the hyperactive mutant achieves a higher PET binding affinity through both a faster on-rate and a slower off-rate. Finally, the landing rate for the Qdot control lacking PETase was 394 ± 113 / μm^2^ · s · M (N = 11), roughly 1/3 of wild-type PETase (Fig. 3D, dashed line). This nonspecific Qdot-PET interaction in the absence of enzyme means that the observed landing rates of the three enzymes are somewhat overestimated; it follows that the measured enhancement of the landing rate of the hyperactive mutant is a lower limit of the effect. A rough correction for this nonspecific binding fraction is to consider the dashed red line in Fig 3D as the baseline.

### pNPA acts as a competitive inhibitor for PET binding

To more thoroughly confirm that the observed binding interactions truly reflect the enzymatic activity of the wild-type and mutant PETases, we used a PETase inhibitor to more clearly separate out specific binding from the various non-specific interactions that may be occurring. 4-Nitrophenyl acetate (pNPA) is a model soluble substrate that is used for measuring the activity of esterases including PETases (7). We reasoned that when used at high concentrations, pNPA should serve as a competitive inhibitor of the PETases, occupying the active site and blocking enzyme binding to the immobilized PET film. Thus, we repeated the single-molecule binding experiments in the presence of 4 mM pNPA.

pNPA reduced the binding duration of wild-type PETase and the hyperactive variant, but had negligible effect on the binding durations of the impaired mutant and the control Qdots lacking bound PETase (Fig. 4A). The clear effects of pNPA on the wild-type and hyperactive PETase provide further validation that the fast binding durations are reporting productive interactions of the enzymes with the immobilized PET. In contrast, the finding that the binding duration of the impaired mutant did not change in the presence of pNPA calls into question whether this mutant is specifically binding to the immobilized PET at all under our conditions.

**Figure 4.**
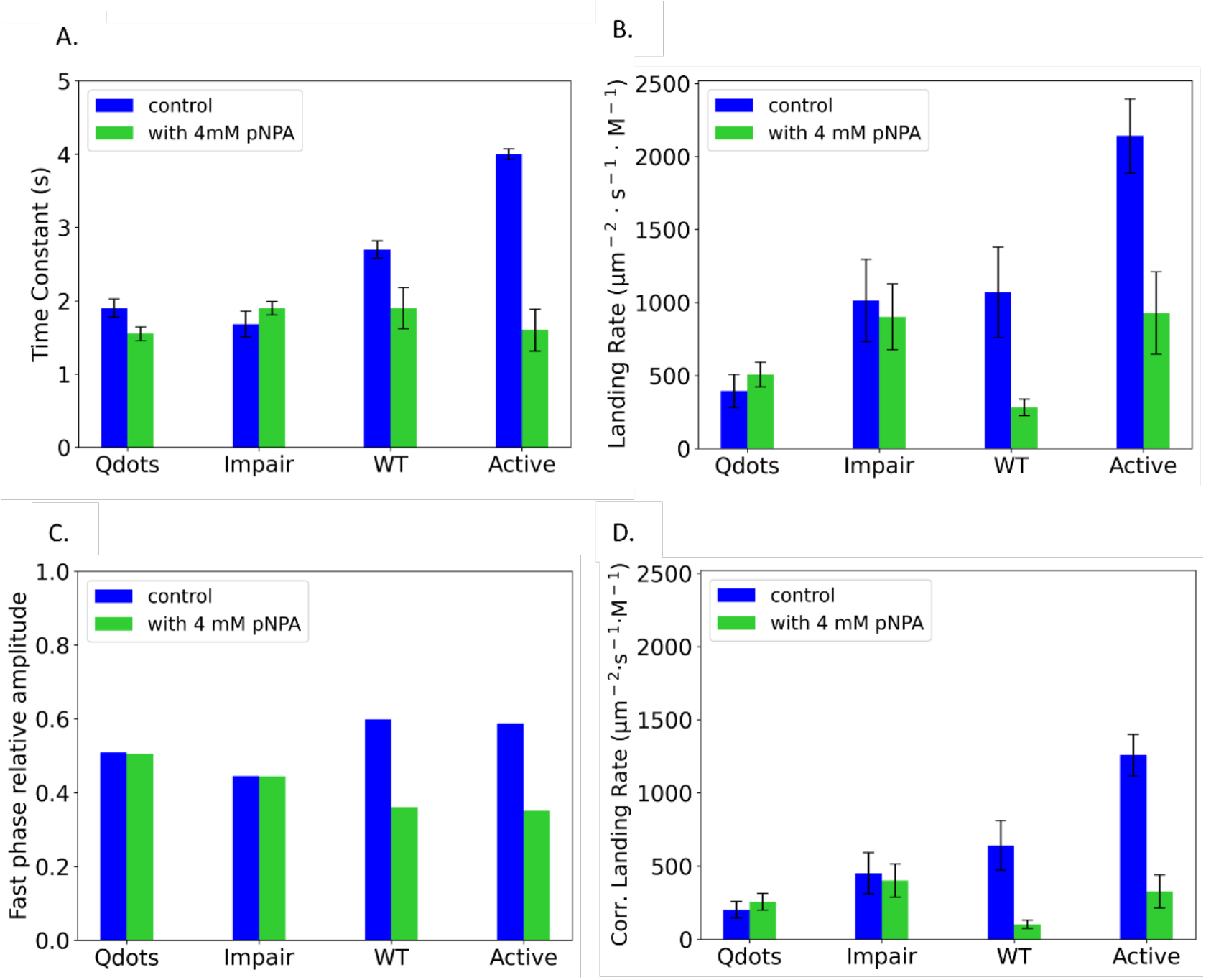
The competitive inhibitor pNPA blocks PETase binding to PET. (A) Time constant for fast phase of Qdot control, impaired mutant, wild-type PETase, and hyperactive mutant in the presence (green bar) or absence of 4 mM pNPA (blue bar). See Fig. S2 for binding duration distributions. (B) Overall landing rate of Qdot control, impaired mutant, wild-type PETase, and hyperactive mutant in the presence (green bar) and absence of 4 mM pNPA (blue bar). (C) Relative amplitude in biexponential fit of the fast phase for Qdot control, impaired mutant, wild-type PETase and hyperactive mutant in the absence (blue bar) and presence of 4mM pNPA (green bar). (D) Corrected landing rate (defined as the overall landing rate multiplied by the relative amplitude of the fast phase) for Qdot control, impaired mutant, wild-type PETase and hyperactive mutant in the presence (green bar) and absence of 4 mM pNPA (blue bar). Error bars are SE of the fit.

The impacts of pNPA on the landing rate mirrors the effects on the binding duration. Specifically, pNPA strongly diminished the landing rate of the wild-type and hyperactive mutant, whereas it had a negligible effect on the landing rate of the impaired mutant and the Qdot control (Fig. 4B). To better separate out contributions to the landing rate from specific binding versus nonspecific binding, we first quantified the relative amplitudes of the fast and slow phases from the biexpontial fits (Fig. 4C). For wild-type and the hyperactive mutant, the amplitudes of the fast durations were diminished by pNPA, consistent with the inhibitor blocking the specific (fast) interactions. As expected, the relative amplitudes of the impaired mutant and the Qdot control were unaffected by pNPA. In the final analysis in Fig. 4D, we multiplied the overall landing rate from Fig. 4B by the relative amplitude of the fast phase from Fig. 4C. This corrected landing rate corresponds to the degree to which the pNPA inhibitor blocks functional binding of the PETases to the immobilized PET. In summary, the pNPA results argue that both wild-type and hyperactive mutant PETase bind specifically and reversibly to the PET, with hyperactive mutant having a faster relative on-rate.

## Discussion

Despite a number of studies that have used bulk assays and computational simulations to investigate the enzymatic activities of PETases, there has been little work at the single-molecule level aimed at uncovering their enzymology. Because the single-molecule approaches developed here enable us to measure and quantify binding events directly, they provide new information that can be integrated into more comprehensive studies of the enzymology of PETases and similar enzymes. These molecular insights can point to new directions for optimizing PETases to the point that they can be scaled up to tackle the significant problem of microplastics pollution in the environment.

The most challenging aspect of these single-molecule investigations was separating specific binding of the enzymes from nonspecific binding by the Qdot probes. Due to autofluorescence of the PET sample, labeling the PETases with standard fluorophores such as rhodamine resulted in an insufficient signal-to-noise for reliable detection. Instead, we turned to Qdots, which gave a very good signal-to-noise, but suffered from non-specific binding to the PET. We explored using surfactants to reduce nonspecific binding but were unable to find conditions that eliminated nonspecific binding while retaining specific binding. Thus, we carried out a series of control experiments to determine source of the fast and slow time constants we identified in our biexponential dwell times fits.

By characterizing Qdots lacking bound PETase and varying laser intensity to characterize photobleaching, we concluded that the slow time constant (∼tens of seconds) of our biexponential fit to the dwell time distributions was due to nonspecific binding. The Qdot control lacking any bound enzymes had a residual binding interaction with a dwell time of 1.9 ± 0.12 s, but the landing frequency was substantially lower than Qdots containing a bound PETase (Fig. 3C and D). This lower landing rate and the elevated binding durations of the wild-type PETase and the hyperactive mutant were the first indication that the assays were detecting specific PET-PETase interactions. The strongest evidence that the fast binding population was indeed specific binding of the PETase to the immobilized PET was the fact that in the presence of the inhibitor pNPA, both the dwell times and the landing rates of the wild-type PETase and the hyperactive mutant fell to near the Qdot control (Fig. 4). The most important experimental result was that the mean dwell time of wild-type PETase was 2.7 s, the dwell time of the hyperactive S238F/W159H mutant was 4.0 s, and the dwell time of the impaired W185A mutant was 1.7 s, which was similar to control Qdots.

What can single-molecule imaging of enzymes tell us about their kinetic parameters? The Michaelis-Menten framework consists of a reversible binding step, with rates k_on_ and k_off_, and a catalytic step, k_cat_.

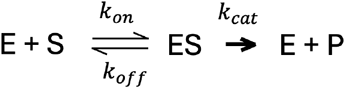

The standard assumption is that product unbinding is fast relative to the catalytic step and is thus lumped into k_cat_. For single-molecule visualization of enzymes binding to an insoluble substrate, we are visualizing the ES complex (meaning the clock starts when the fluorescent enzyme binds to the surface). What we measure is dissociation from the surface, which is described by the apparent dissociation rate,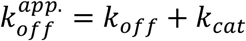 . If we assume that the fast binding durations we measure are reflecting the residence time of the enzyme on the PET substrate, then this apparent off-rate can be calculated from our binding durations as 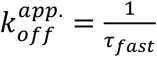 . From the fast time constants in Fig. 3C, 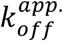 are 0.37 s^-1^ for the wild-type, 0.25 s^-1^ for the hyperactive mutant, and 0.6 s^-1^ for the impaired mutant.

The first result that comes out of this analysis is that the upper limit for k_cat_ for wild-type PETase is 0.37 s^-1^. This conclusion comes from the fact that dissociation of the enzyme from the substrate is either through unbinding before catalysis (k_off_) or via a catalytic event followed by dissociation from the “product” (cleaved PET in our case), with rate k_cat_. It should be noted that the catalytic event may be considerably faster than this, meaning that the lumped k_cat_ parameter is rate-limited by product release. However, the functional turnover rate would be the same in this case, as would the apparent k_cat_ that we observed. This relatively slow turnover rate presumably is related to the crystallinity of PET film and the fact that experiments were performed at room temperature. Previous work showed that higher degrees of crystallinity lead to greater resistance of PET to enzymatic hydrolysis (17, 18). It is proposed that as temperatures approach the glass transition temperature of PET, the mobility of the PET chain increases and results in more rapid enzymatic hydrolysis (11, 19). Using bulk assays with the same wild-type PETase as used here, Erickson et al measured a k_cat_ of 1.5 s^-1^ on amorphous PET and 0.8 s^-1^ on crystalline PET at 30 °C (20), somewhat faster, but within the range of our k_cat_ of 0.37 s^-1^ at 22 °C.

The enzyme dissociation rates that we measure also provide insights into how mutations alter the kinetic parameters of the enzymes. We interpret our data in the context of the simple Michaelis-Menten model showin in Fig. 5. The hyperactive mutant, S238F/W159H was chosen by Austin *et al*. because the mutations narrowed the active cleft of the enzyme to match that of other cutinases (6). Induced fit docking simulations predicted that this double mutant would have much higher affinity for PET, though no predictions were made on how the mutations may alter k_cat_. We find that 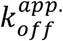 drops from 0.37 s^-1^ for the wild-type to 0.25 s^-1^ for the hyperactive mutant. First, because a faster k_cat_ predicts a faster 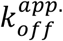 (Fig. 5), our data argue that the greater surface erosion and decrease in crystallinity seen with this mutant in a previous study (6) do not result from an elevated k_cat_. Consistent with this, Erickson et al. measured a k_cat_ of 0.5 s^-1^ for this hyperactive mutant at 30 °C, slower than their wild-type k_cat_ of 0.8 – 1.5 s^-1^ in the same study (20). This value at 30 °C is twice our inferred k_cat_ at 22 °C, which is reasonable agreement. However, this lower k_cat_ measured by Erickson et al. seems at odds with the elevated PET surface erosion of this mutant (6). One possible explanation of our slower apparent off-rate is that the rate of reversible binding from the substrate, k_off_, is slowed in the hyperactive mutant. The first effect of slowing k_off_ is a lower K_M_, which will improve performance at sub-saturating substrate concentrations. The second effect is that for every enzyme-substrate encounter, there is a higher probability that the enzyme will hydrolyze the substrate (with 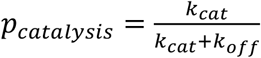Fig. 5) rather than simply unbinding without acting on the substrate (with probability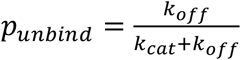).

**Figure 5.**
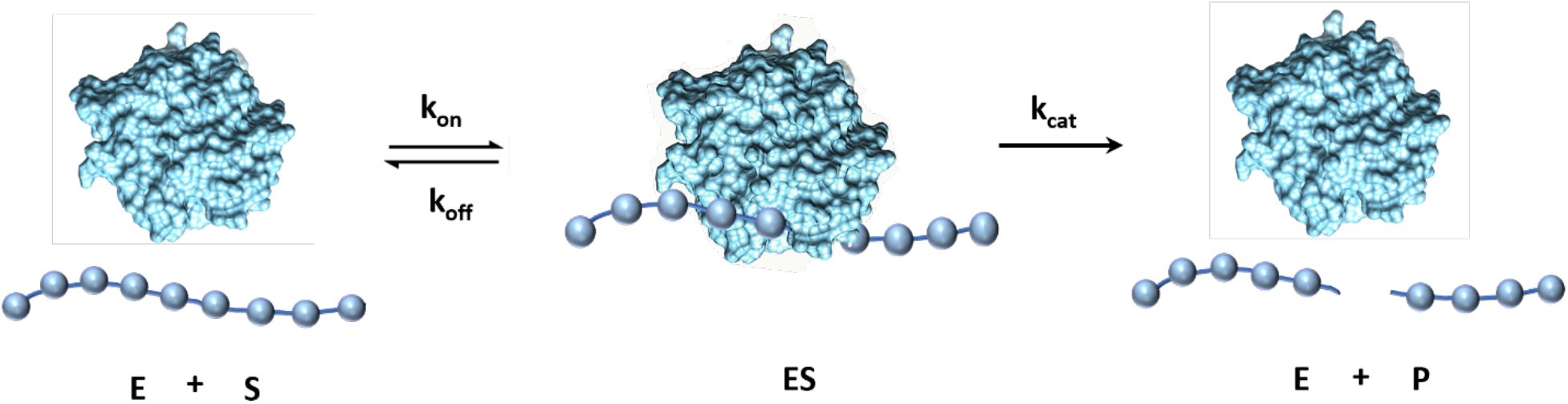
Kinetic model of PETase binding to PET film. Free PETase in solution binds to PET film reversibly with association rate constant k_on_ to form an enzyme-substrate complex. The complex then either disassociates with rate constant k_off_ or proceed to the PET cleavage step with turnover rate k_cat_, followed by rapid dissociation. Qdot-labeled PETase molecules bind in the ES state and dissociate with an apparent rate constant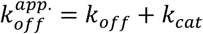, which is equal to the inverse of the fast phase dwell time. For each binding interaction, the probability that catalysis proceeds rather than the enzyme unproductively dissociating is: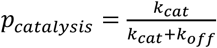 .

We can also use this analysis to interpret the impaired mutant. Previous work suggested that residue W185 reorients upon productive binding to PET and may contribute *π*-stacking interactions with aromatic groups in the PET substrate (6). The W185A mutation was found to have reduced surface erosion activity on PET and a smaller reduction in PET crystallinity, compared to wild-type (6). First, it should be noted that, because we found that τ_*fast*_ for the impaired mutant was indistinguishable from that for Qdots alone (Fig. 3C), the 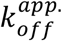 of 0.6 s^-1^ for the impaired mutant should be considered a lower limit. Importantly, the landing rate of the impaired mutant was considerably faster than Qdots alone (Fig. 3D), which argues that the impaired mutant is not totally inactive. The simplest interpretation of the faster apparent off-rate is that the off-rate of the impaired mutant from the PET substrate is considerably faster than wild-type. This effect makes sense based on the proposed role of W185 in PET binding, and in bulk assays would lead to a larger K_M_. Another way to describe the mutation is that for every binding encounter, the probability of a catalytic event, 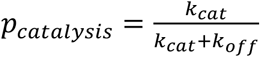, is smaller, making the enzyme less productive.

## Conclusion and future outlook

We show here that single-molecule imaging can be used to deduce kinetic parameters of PETase enzymes. This approach provides a new tool that can be combined with existing bulk methods and computational tools to engineer more active and robust PETases that address microplastics pollution. Further development of this single-molecule approach will require reducing the non-specific adsorption that complicates interpretation of the kinetic data. Potential solutions include using probes other than Qdots to label the enzymes, improved processing of the PET substrate to reduce the autofluorescence, and identifying surfactants or other alterations in buffer conditions that minimize nonspecific binding. Notably, more active enzymes that have higher k_cat_ values are predicted to bind for shorter durations, and nonspecific Qdot binding currently limits the upper limit of apparent k_off_ values that can be measured. One approach that has been explored for improving PETase function is attaching a Carbohydrate Binding Module (CBM) or another hydrophobic domain to PETases to improve their PET binding affinities (21, 22). If such a strategy is successful in creating a processive enzyme that diffuses along the surface while carrying out multiple chain cleavage reactions, this activity should be measurable as both a slower apparent off-rate of the enzyme measured by the current approach, as well as diffusive movements that could be detected by single-molecule tracking. In conclusion, the methods developed here provide a new tool for screening engineered PETases with enhanced functions, which should contribute to multidisciplinary efforts to use biocatalysts to reduce microplastics pollution in the environment.

## Acknowledgements

The authors thank members of the Hancock lab for helpful suggestions and Dr. Justin Brown for advice in PET preparation. This work was supported by NIH grant R35GM139568.

## Supporting Information

**Figure S1.**
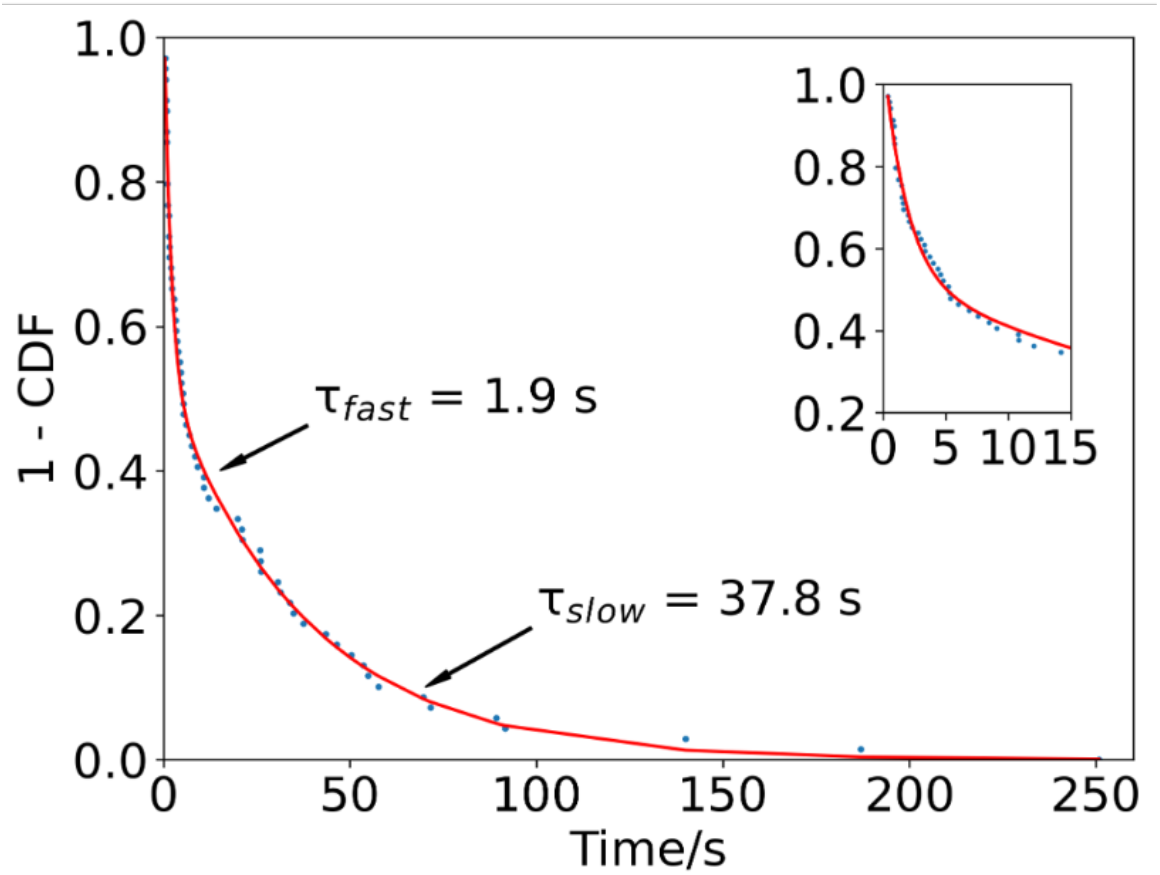
(related to Fig. 3): Binding durations of PETase-free Qdot control. Distribution of binding durations of Qdot control, fit by a biexponential. Inset: first 15 s to show details of fast phase.

**Figure S2.**
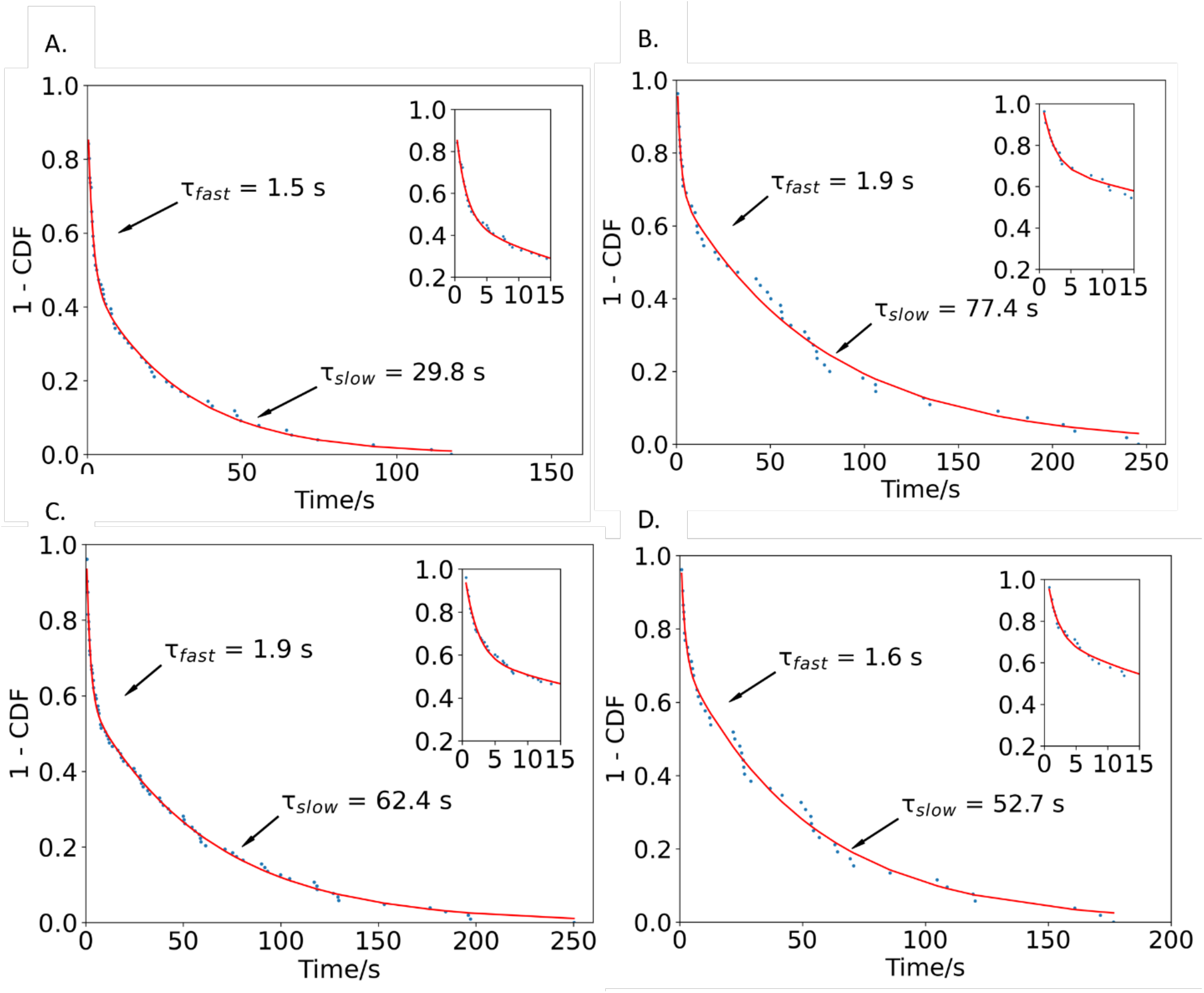
(related to Fig. 4): Binding durations in the presence of 4 mM pNPA. (A) Distribution of binding durations of Qdot control in the presence of 4 mM pNPA, fit by a biexponential. Inset: first 15 s to show details of fast phase. (B) Distribution of binding durations of wild-type PETase in the presence of 4 mM pNPA, fit by a biexponential. Inset: first 15 s to show details of fast phase. (C) Distribution of binding durations of the inactive PETase mutant in the presence of 4 mM pNPA, fit by a biexponential. Inset: first 15 s to show details of fast phase. (D) Distribution of binding durations of the hyperactive PETase mutant in the presence of 4 mM pNPA, fit by a biexponential. Inset: first 15 s to show details of fast phase.

## Notes

### Competing Interest Statement

The authors have declared no competing interest.

### Summary of Updates

Manuscript has been reformatted, discussion was further developed, and references were added and corrected.

